# TSSK homologue regulates the expression of *protamine* and *mosquito testes specific* genes in *Anopheles stephensi*

**DOI:** 10.1101/2024.11.14.623542

**Authors:** Keshav Kumar Meghwanshi, Chhavi Choudhary, Pooja Rohilla, Rajnikant Dixit, Vishal Kumar Saxena, Jayendra Nath Shukla

## Abstract

The testis-specific serine/threonine-protein kinase (*tssk*) gene is known to play an important role in spermiogenesis in mammals. This study identifies *tssk* homologues, *As_tssk3* and *Aea_tssk1* in *Anopheles stephensi* and *Aedes aegypti*, respectively. Both *As_tssk3* and *Aea_tssk1* were found to express in a male-specific manner throughout the development. Efficient knockdown in the expression of *As_tssk3* gene was observed upon feeding *An. stephensi* larvae with target dsRNA. *As_tssk3* knockdown led to a reduced sperm reservoir in male adults. Additionally, the knockdown of *As_tssk3* resulted in the reduced expression of two male specifically expressed genes, *protamine & mosquito testes specific* (*mts-* a sperm-specific marker) homologues of *An. stephensi*. *In-silico* interaction analysis suggests that As_TSSK3 protein interacts with the Nucleosome Assembly Protein1 like-4 (NAP1like-4), a putative transcription factor. We hypothesize that As_TSSK3 regulates the transcription of *protamine* and *mts* genes via NAP, as both NAP and *protamine* have previously been shown to function in the establishment and maintenance of chromatin dynamics, in independent studies. The study presented here is the first report of characterisation of *tssk* homologues in any mosquito species. The knockout studies and phosphorylation assays would provide more insights to explore the mechanism by which *As_tssk3* regulates *protamine* and *mts* genes in *Anopheles stephensi*.

**Highlights:**

- This study identifies the *tssk* homologues in *Ae. aegypti* and *An. stephensi* and showed their male-specific expression in both the mosquito species.
- dsRNA mediated knockdown of *As_TSSK3* resulted into the males with a reduced size of sperm reservoir.
- dsRNA mediated knockdown of *As_TSSK3* led to the reduction in the expression of *As_prot* and *As_mts*, homologues of protamine and mosquito testis specific, genes in *An. stephensi*.
- In-silico studies suggest that As_TSSK3 regulates the expression of *As_prot* and *As_mts* genes via NAP.

## 2. INTRODUCTION

Spermatogenesis, a fundamental reproductive process, involves a series of molecular and biochemical steps (Neto et al., 2016). The proteins synthesized during the process of spermiogenesis play a crucial role in the maturation and activation of spermatozoa. In addition, several post-translational modifications (PTMs) involving ubiquitination, acetylation, SUMOylation and phosphorylation have also been reported to play an important role in sperm cell development. A broad group of kinases including MAPKs, PLKs, AURKC, and the TSSK family of kinases have been shown to be essential for spermatogenesis and male fertility (Ben khelifa et al., 2011; Bettencourt-Dias et al., 2005; Carmena et al., 1998; Jordan et al., 2012; M. W. M. Li et al., 2009). The Testis-Specific Serine/Threonine Kinase (TSSK) family is a conserved group of protein kinases which are primarily expressed in the testes and regulate spermatogenesis (Salicioni et al., 2020a). Members of the TSSK family, characterized by the presence of a serine/threonine protein kinase catalytic (S-TKc) domain, were first identified in mice (Bielke et al., 1994). This kinase family includes various members, for example TSSK-1 to TSSK-6 and are involved in regulating different aspects of spermatogenesis. Besides rodents, TSSK homologues have also been identified in other mammals (Kueng et al., 1997), molluscs (Kim et al., 2019), birds (Słowińska et al., 2020), and insects (X. Yang, 2024).

A recent study has led to the identification of TSSK homologue (*dtssk*) in *Drosophila melanogaster* (Zhang et al., 2023). *dtssk* is specifically expressed in the testis of adult male flies, and its mutation causes male sterility. *dtssk* mutant flies exhibit several distinct defects in their spermatids, including aberrant nuclear morphology, disorganized flagella, and impaired DNA condensation. These abnormalities arise because of impairment in the histone-to-protamine transition during spermiogenesis, a process essential for proper sperm maturation. Further, phosphoproteomic analysis led to the identification of multiple substrates of dTSSK protein, many of which are involved in microtubule dynamics, spermatid differentiation, and development. These results provide valuable insights into the molecular mechanisms behind male infertility linked to defects in TSSK. Beyond *Drosophila*, TSSK homologues have also been identified in several other insects, including the oriental fruit fly, *Bactrocera dorsalis* (Sohail et al., 2019); melon fly, *Zeugodacus cucurbitae* (Zhai et al., 2023) and codling moth, *Cydia pomonella* (X. Yang, 2024). In all these species, TSSK homologues are predominantly expressed in the testes and are crucial for male fertility.

dsRNA mediated knockdown of TSSK homologues, *tssk1* in *B. dorsalis* and *Zctssk* in *Z. cucurbitae*, resulted in a significant reduction in sperm count in these insects. Besides, a reduced hatching rate was also recorded in the eggs laid by virgin females mated with TSSK knockdown males (Ashok et al., 2024; Sohail et al., 2019; Zhai et al., 2023). In *C. pomonella*, a total of 5 TSSK (TSSK1, TSSK1a, TSSK2, TSSK2a, & TSSK4) homologues were identified, with TSSK4 showing high expressions in testis. CRISPR mediated knockouts of each of the five *tssk* genes independently resulted in decreased sperm motility, reduced sperm count, and abnormal development of eggs, leading to lower hatching rates. Additionally, TSSK2, TSSK2a and TSSK4 knockouts were completely sterile.

*Aedes & Anopheles* mosquitoes, belonging to the order Diptera, are the vectors of several parasites and viruses. However, there is no study related to TSSK, and its homologues have been performed in any of the mosquito species till date. In this study, we identified the TSSK homologues, *As_tssk3-like* (*As_tssk3*) and *Aea_tssk1-like* (*Aea_tssk1*) in *Anopheles stephensi* and *Aedes aegypti*, respectively. Developmental profiling revealed that, both the genes (*As_tssk3* and *Aea_tssk1*) exhibited male-specific expression during pupal and adult stages. Though the expression of *As_tssk3* was observed in head, gut and gonadal tissues of male adults, its expression was significantly higher in gonads (testes and accessory glands). Feeding dsRNA targeting *As_tssk3* to 3^rd^ instar larvae led to a significant reduction in its expression in *An. stephensi* males. These males possessed a smaller sperm reservoir as compared to that in control males.

Interestingly, the knockdown of *As_tssk3* also resulted in a reduced expression of *protamine* homologue (*As_prot*) and a sperm marker gene (*mosquito testis specific; As_mts*) (Krzywinska & Krzywinski, 2009; Kumari et al., 2022), compared to that in control males. We hypothesized that a transcription factor targeted by As_TSSK3 protein may regulate the expression of *As_prot* & *As_mts* genes. To explore this, we searched for transcription factors involved in spermatid development and differentiation among dTSSK targets. As a result, we identified Nucleosome Assembly Protein (NAP) as a key target of dTSSK. NAP is known for its crucial role in histone mobility and transcriptional regulation in several eukaryotic organisms was found to be one of the targets of dTSSK (Gill et al., 2022; Park & Luger, 2006b, 2006a). Further, we identified the homologue of NAP, As_NAP1 (As_NAP1 like-4), a 341 amino acid protein in the *An. stephensi* database, which possesses multiple phosphorylation sites suggesting it to be the potential target of As_TSSK3. Additionally, our *in-silico* interaction studies suggested a strong interaction between As_TSSK3 and the As_NAP1 of *An. stephensi*.

Overall, this study identifies TSSK homologues in *An. stephensi* and *Ae. aegypti* and investigates the potential role of *As_tssk3* in spermatogenesis in *An. stephensi*. It also explores the pathway through which As_TSSK3 may regulate *As_prot*, and *As_mts* genes, highlighting a transcriptional association of As_TSSK3 with *As_prot* and *As_mts* regulation. Additionally, the sex-specific role of *As_tssk3* in maintaining the sperm reservoir highlights its potential as a target for unraveling the molecular mechanisms of spermiogenesis in mosquitoes.

## 3. Material and Methods

### 3.1 Mosquito rearing

The *Anopheles stephensi* and *Aedes aegypti* cultures were maintained at 26°C ± 2°C, with 70–80% relative humidity and a 12:12 day/night cycle as described previously (Kumar et al., 2021; Kumari et al., 2022). Sexing of pupae and adults were done based on sexually dimorphic characters described earlier (Boo, 1980; Yamany et al., 2024).

### 3.2 TSSK homologues in Aedes aegypti and Anopheles stephensi

Protein sequences of TSSK homologues (XP_021705955.1 and XP_035919898.1) of *Ae. aegypti* and *An. stephensi*, respectively were retrieved from the NCBI database. These protein sequences were used to perform BLAST (tBLASTn) analysis in the Expressed Sequence Tags (EST) databases of *Ae. aegypti* and *An. stephensi* which led to the identification of corresponding mRNA sequences, XM_021850263.1 and XM_036064005.1, respectively. These mRNA sequences were aligned with their corresponding genomic contigs (NIGP01000003.1, JABEOT010000002.1) using NCBI splign tool to deduce the exon-intron boundaries (https://www.ncbi.nlm.nih.gov/sutils/splign/splign_note.html). The open reading frames (ORFs) in the mRNA sequences of these genes were determined using the Expasy Translate tool (https://web.expasy.org/translate/). Domains present in these TSSK protein homologs were identified using Conserved Domain Database (CDD) of NCBI (https://www.ncbi.nlm.nih.gov/Structure/cdd/wrpsb.cgi). Supplementary table 1 (S1) shows the accession IDs of transcript and genomic sequences of all the genes analysed in this study.

### 3.3 RNA isolation, cDNA synthesis and RT-PCR

Total RNA was extracted from different developmental stages including eggs, larval instars, male and female pupae, and male and female adults of *An. stephensi* and *Aedes aegypti* using Trizol reagent (Invitrogen). Total RNA was also isolated from the tissues (head, gut, & gonad) of male adults of *An. stephensi*.

RNA samples were DNase (Promega) treated as per the manufacturer’s protocol to remove any residual genomic DNA. The concentration of RNA samples was taken using microvolume spectrophotometer (NanoDrop 2000, Thermo Scientific). One microgram (µg) of DNase treated RNA sample was used to synthesize cDNA using GoScript Reverse Transcriptase (Promega) following the manufacturer’s instructions. Subsequently, cDNA samples were diluted 2-fold with nuclease free water (Ambion) for their use in PCR reactions.

Gene-specific primers were designed using Primer-3 software (Primer3 Input, version 0.4.0). Tissue and stage-specific cDNAs were used as templates to perform PCR using gene-specific primers (Table S2) and GoTaq PCR master mix (Promega). All the PCR reactions were performed in Proflex 32 thermocycler (Applied Biosystems) with the following cycling parameters; an initial denaturation at 94°C for 2 minutes, followed by 32 cycles of denaturation at 94°C for 30 seconds, annealing (primer-specific, Table S2) for 30 seconds, extension at 72°C for 30 seconds, and a final extension at 72°C for 10 minutes. PCR amplified products were electrophoresed on 1.5% agarose gels and visualized using a gel documentation system (Bio-Rad).

### 3.4 qPCR and statistical analysis

Quantitative PCR (qPCR) was performed to analyse the relative mRNA expressions, using cDNA as template along with gene-specific primers (Table S2) and Luna SYBR Green qPCR master mix (NEB), following the manufacturer’s protocol. The *An. stephensi actin* gene was used as an endogenous control for the normalization of the expression data (Collins et al., 2014) and, the gene expression levels were analyzed by 2^^ΔΔCt^ method (Livak & Schmittgen, 2001). The qPCR cycling parameters included an Initial denaturation at 95°C for 5 minutes, followed by 40 cycles of denaturation at 95°C for 10 seconds, annealing at 53°C for 30 seconds, extension at 72°C for 30 seconds, and a final extension at 72°C for 10 minutes. Melt curve analysis was carried out in a temperature range of 60°C-95°C with an increment of 0.5°C per 5 seconds, to rule out the presence of any non-specific amplification.

The statistical analysis was performed using GraphPad Prism (9.5.1). Gene expression levels and the knockdown efficiency of *As_tssk3* were performed using t-test analysis. One-way ANOVA was used to compare *As_tssk3* expression levels between different developmental stages and between different tissues of *An. stephensi*. Three biological replicates per group were used in the study, and all the experiments were performed in triplicates.

### 3.5 dsRNA synthesis, droplet feeding assay and mating experiments

A region of *As_tssk3* (Fig. 1C) was amplified by PCR using cDNA as template and gene-specific primers having T7 promoter sequence (TAATACGACTCACTATAG) at their 5’end. Purified amplicon was then used as a template in *in-vitro* transcription reaction to synthesize dsRNA using Megascript T7 transcription kit (Invitrogen). Similarly, the dsRNA corresponding to *gfp* gene was also prepared for use as a control.

**Fig. 1.**
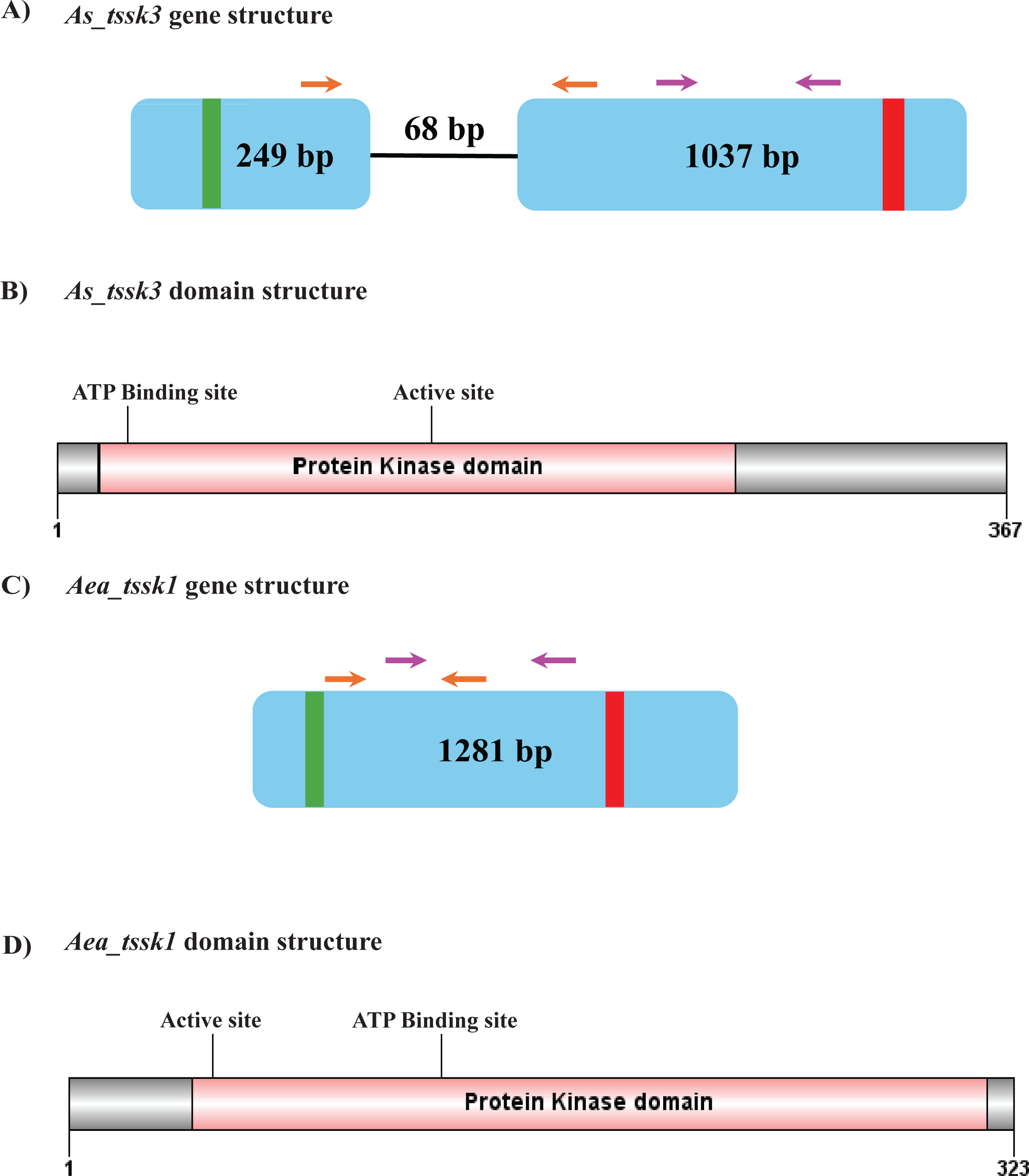
Schematics showing the gene structures and domains present in *As_tssk3* and *Aea_tssk1* genes. **A and C** represent the genes structures of *As_tssk3* and *Aea_tssk1* genes. The *As_tssk3* gene has two exons and one intron whereas *Aea_tssk1* is a mono exonic gene. Vertical lines represent the start (green) and stop (red) codon sites. Arrows represent the location of forward and reverse primers (Orange-qRT primers and purple dsRNA primers) in the genes. **B and D** represent the relative positions of domains present in As_TSSK3 & Aea_TSSK1 proteins.

*As_tssk3* or control dsRNA was fed to two different groups of 3^rd^ instar larvae for 2 hours using droplet feeding method (Rodrigues et al., 2017; J. Yang & Han, 2014). For this, a group of larvae (N=15) were placed in a small container having 5 µg of dsRNA in 75µl water nuclease free water. A total of 10 such groups were maintained for feeding of both control and target gene-specific dsRNA. After 2 hours, larvae from each group of dsRNA were mixed and placed in a separate water tray under normal rearing conditions. Males were separated from females after pupation with the help of sexually dimorphic structure (Leite et al., 2024; SUGUNA et al., 1994). Male pupae and adults were collected for RNA isolation, which was used to assess the gene silencing efficiency using qPCR.

#### 3.5.1 Oviposition assay

After sex sorting of knockdown pupae, male pupae were maintained in separate rearing cages (control and target groups) until the adult emergence. After 48hrs, knockdown male adults were mated with control virgin female adults. Subsequently, these females were blood-fed, and fully engorged females were selected for egg laying. The selected females were placed in individual cups (2 females per cup), with water and blotting paper attached to the walls. These blotting papers were collected after over-night egg laying and the number of eggs laid was counted manually (Kumari et al., 2022).

### 3.6 In-silico studies

The interaction of As_TSSK3 with other proteins (Table S5) was studied using STRING analysis (Szklarczyk et al., 2023). The domain structure of As_TSSK3 and phosphorylation sites in the As_NAP1 protein were predicted using GPS 5.0 software (Xue et al., 2008). The protein sequences related to As_TSSK3 were retrieved from NCBI database followed by phylogenetic analysis using Mega 11 software (Tamura et al., 2021).

### 3.7 Microscopic assay

To analyze the morphological changes in the gonad of *As_tssk3* knockdown males, control and knockdown males were cold anesthetized and dissected under microscope (Nikon, C-LEDS). Further, the gonads (testes and accessory glands) of dissected mosquitoes were pulled out using sharp fine needles. Separated tissues were visualized under microscope (Magnus, MLX-TR) and were photographed with the help of a camera attached to the microscope.

## 4. Results

### 4.1 *tssk* homologues express exclusively in male mosquitoes

*As_tssk3* (>XM_036064005.1) gene consists of two exons and one intron, whereas *Aea_tssk1* (>XM_021850263.1) is a mono-exonic gene (Fig. 1A & 1C). *As_tssk3* has ORF of 1104 bp while *Aea_tssk1* has the ORF of 972 bp. *As_tssk3* gene was found to express throughout all the developmental stages (egg, larvae, pupa and adult) of *An. Stephensi* but its expression was significantly higher in male pupae and male adults (Fig. 2A). Although the expression of *As_tssk3* was detected in all the male tissues analysed (head, gut and gonad), its expression was notably higher in gonad (Fig. 2B). Similarly, *Aea_tssk1* expresses during all the developmental stages (3^rd^ instar and 4^th^ instar larvae, sex-specific pupae and sex-specific adult) the expression was limited to males during pupal and adult stages (Fig. S1).

**Fig. 2.**
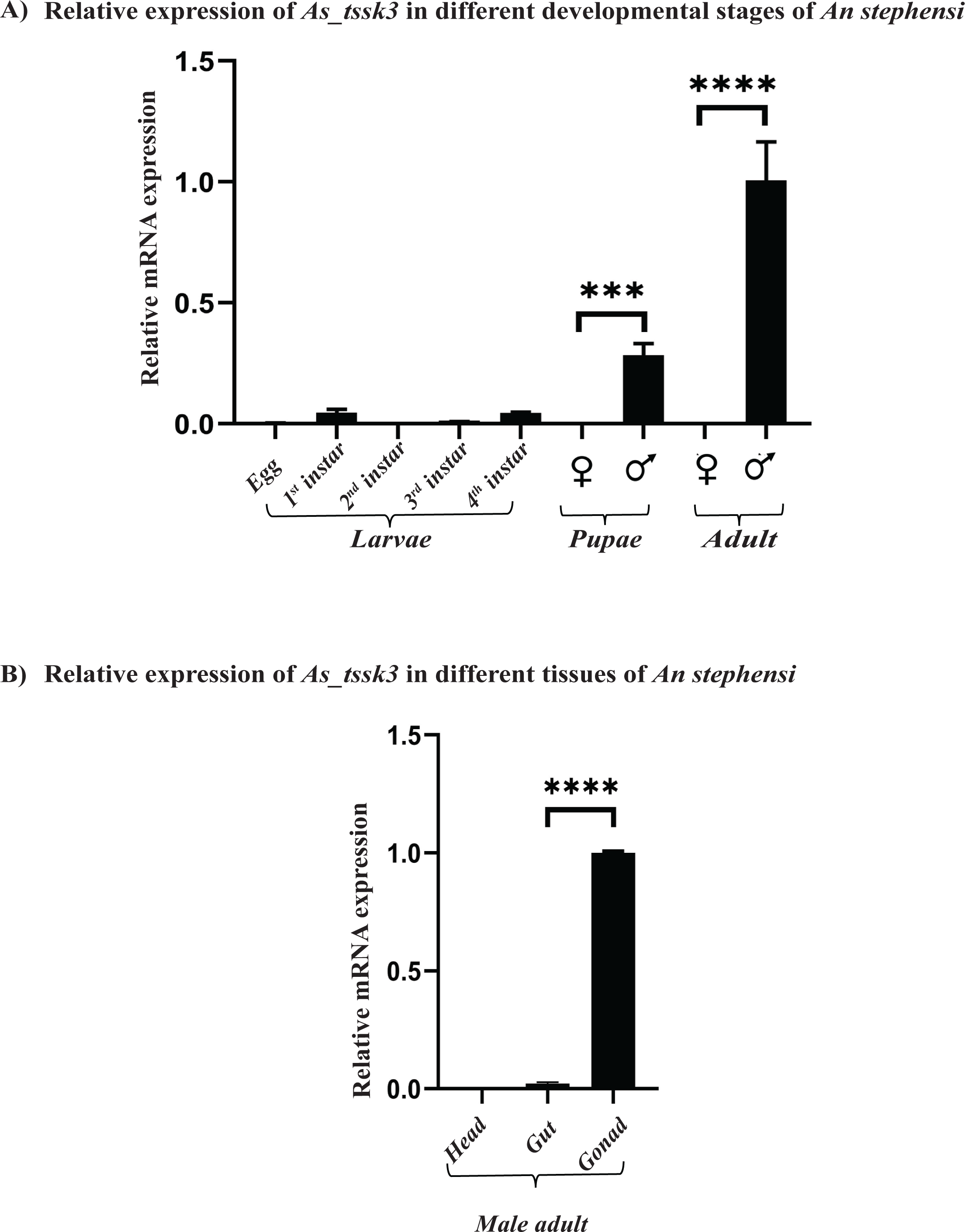
Graphical representation of *As_tssk3* expression in *An. stephensi.* **A)** Relative mRNA expression levels of *As_tssk3* during the developmental stages of *An. stephensi* (****p-value≤0.0001; ***p-value≤0.0008). **B)** Relative mRNA expression levels of *As_tssk3* in different tissues of male adults (****p-value≤0.0001). The *actin* gene *(As_actin)* of *An*. *stephensi* was used as endogenous control. Three biological replicates were used in the experiment and each set was a pool of five individuals or tissues isolated from five male adults for A and B, respectively.

### 4.2 Knockdown of *As_tssk3* reduces the expression of *As_prot and As_mts genes*

Feeding *An. stephensi* larvae with dsRNA targeting *As_tssk3* resulted in a significant reduction in its expression in both male pupae and male adults as compared to that in corresponding controls (Fig.3A & 3B). Additionally, the male adults eclosed from *As_tssk3* dsRNA fed larvae showed a reduced expression of *As_prot* and *As_mts* genes as compared to that in male adults eclosed from control dsRNA fed larvae (Fig.3C & 3D).

**Fig. 3.**
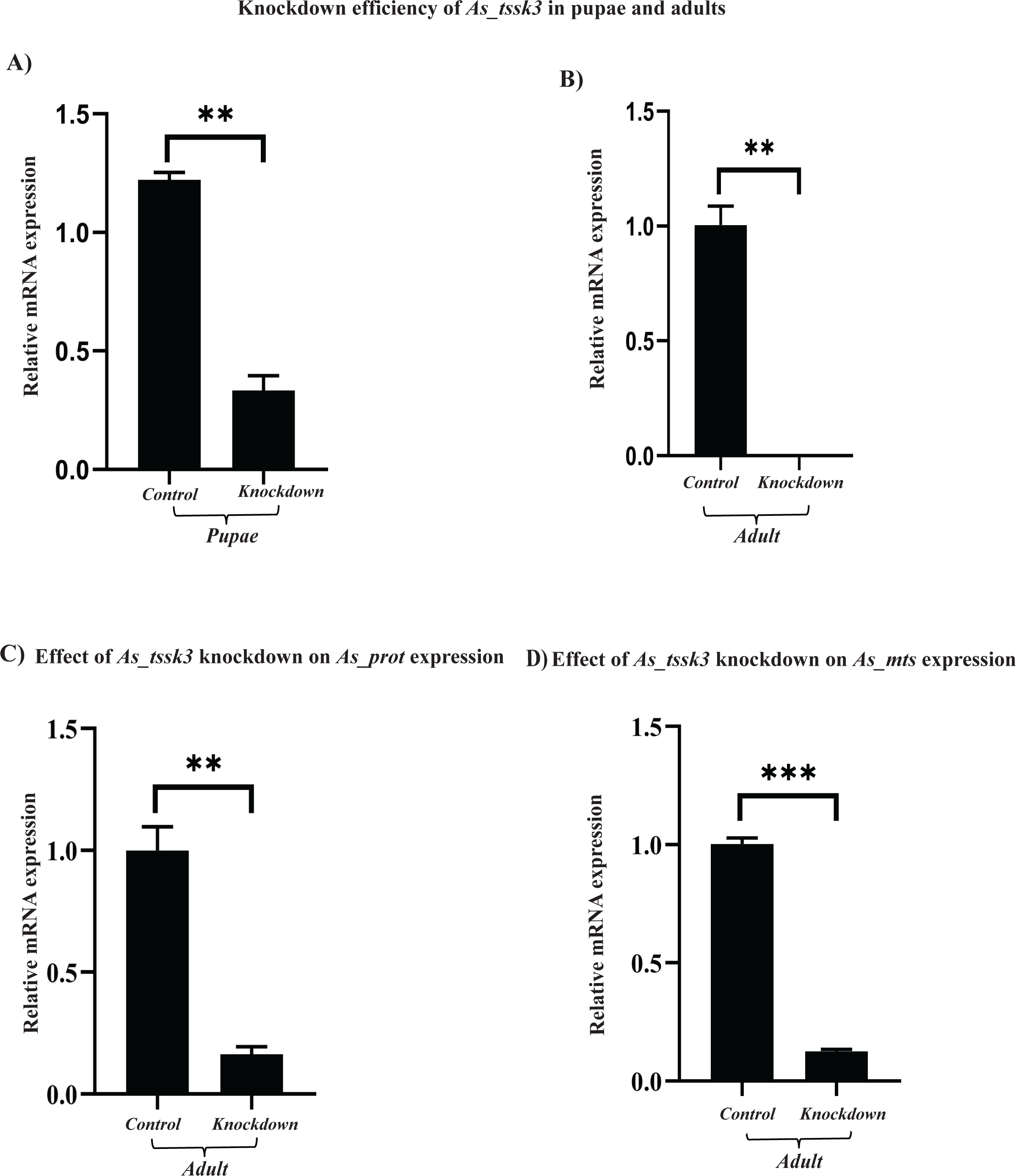
Graphical representation of the effect of dsRNA feeding on the expression of *As_tssk3* in pupae (**A**) and adults (**B**) of *An. stephensi* (**p value ≤ 0.0031; 0.0034, respectively). Graphs showing the effect of *As_tssk3* knockdown on the expression of *As_prot* (**C**) and *As_mts* (**D**) in male adults (**p value ≤ 0.0074; ***p value ≤ 0.0005, respectively). *As_actin* was used as endogenous control. Three biological replicates were used in the experiment and each set was a pool of five individuals.

### 4.3 Reduced sperm reservoir in the testis of *As_tssk3* knockdown male adults

The dissection and analysis of testes from knockdown and control adults revealed reduced sperm reservoir in the testis of some of the knockdown males as compared to the control ones (Fig. 4). The relative sperm reservoir coverage of control testes and knockdown testes was observed as 47% and 34% respectively.

**Fig. 4.**
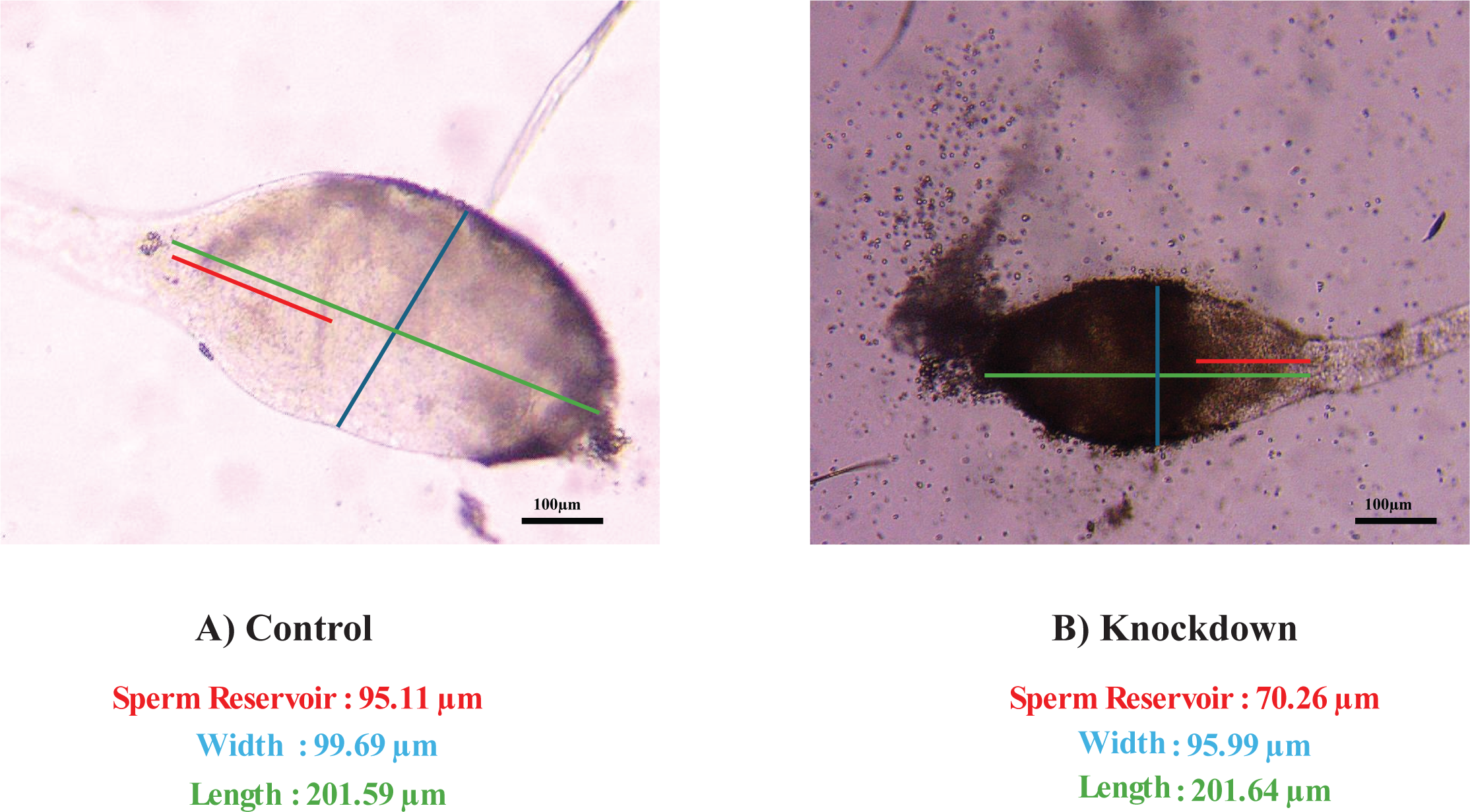
Morphological analysis of the dissected testis from 0-1day old *As_tssk3* control (**A**) and knockdown (**B**) male adults. Colored lines indicate measurements of testes length (Green), width (Blue) and sperm reservoir (Red). Scale bar 100µm. The change in relative sperm reservoir was calculated using the formula: % change formula = Treated - Control / Control X 100.

Specifically, there was a decrease in the size of sperm reservoir by 26%, with knockdown testes exhibiting relative sperm reservoir coverage of 34% as compared to 47% in control testes (Fig. 4 and S3). There was no notable change in the overall size of both control and knockdown testes (Table S3). Furthermore, no significant difference in the number of eggs laid by females mated with *As_tssk3* knockdown males compared to those mated with control males was observed (Table S4).

### 4.4 As_TSSK3 harbors conserved kinase domain and shows interaction with As_NAP1

Similar to other TSSK proteins, As_TSSK3 and Aea_TSSK1 also contain conserved kinase domain, multiple active sites and ATP binding sites (Fig. 1B & 1D). Homologues of nineteen known *dtssk* substrates, which are involved in spermatid development and differentiation, were identified in *An. stephensi* (Table S5). In-silico interaction analysis confirmed the direct interaction of As_TSSK3 with three of these proteins, Nucleosome Assembly Protein1-like4 (As_NAP), Radial Spoke Head (RPH), and Baculoviral IAP repeat-containing protein 6, among which As_NAP showed the maximum interaction (Fig. S2). Further, multiple phosphorylation sites were found to be present in the As_NAP1 proteins (Table S6) suggesting it to be a potential substrate of As_TSSK3.

### 4.5 Evolutionary conservation of As_TSSK3 with other organisms

Phylogenetic analysis using a total of 35 different TSSK proteins from different insect orders, including mosquitoes and outgroups, was conducted to access their evolutionary conservation. The resulting phylogenetic tree indicates that As_TSSK3 shares a common ancestry with TSSKs of *An. gambiae*, *Ae. aegypti* and *Ae. albopictus* species (Fig. 5). The As_TSSK3 shares the closest relationship with the sister clade, *Leptinotarsa decemlineata* (Coleoptera) as both are connected by a common branch. The phylogenetic tree broadly diverged into three main branches showing that species among Coleoptera, Orthoptera, Hymenoptera are closely related, while other groups are distantly diverged.

**Fig. 5.**
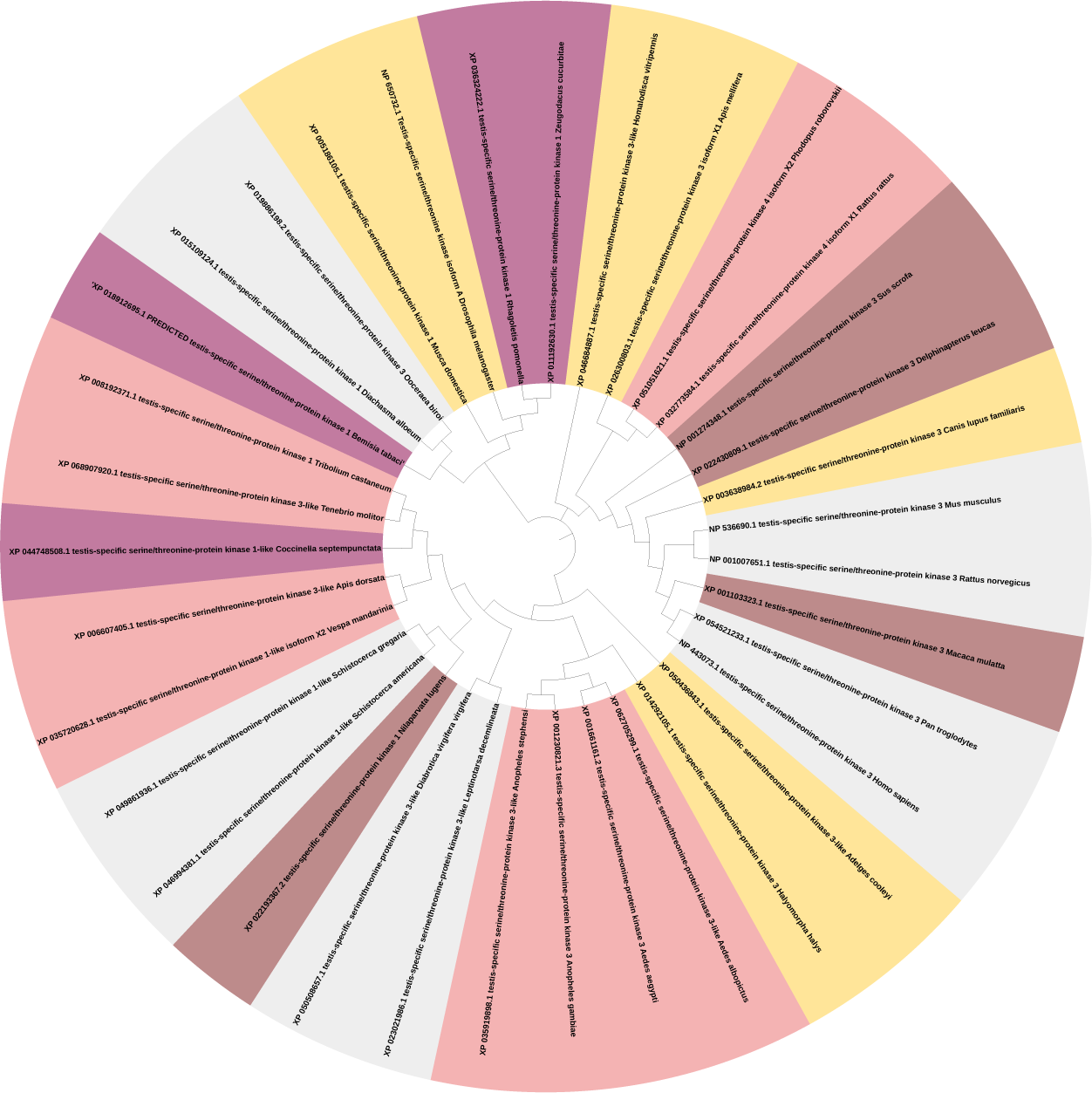
Circular phylogenetic tree of TSSKs homologues from insects and other organisms. Protein sequences were aligned using MEGA11 and the tree was constructed using the neighbour-joining method with a bootstrap value of 1000 replicates.

## 5. Discussion

Spermatogenesis is an essential reproductive process which is governed by complex molecular and biochemical pathways (Brill & Wolfner, 2012; Kanippayoor et al., 2013). Various post-translational modifications (PTMs) facilitated by specific protein kinases are critical to this process and significantly contribute to the maturation of the sperms (Brohi & Huo, 2017; Lewis et al., 2003). Among these, PTMs generated by Testis-Specific Serine/Threonine Kinases (TSSK) have special significance in sperm maturation (Hundertmark et al., 2018; Mao et al., 2022). TSSKs are a part of the CAMK (Calcium/calmodulin-dependent protein kinases) family (Salicioni et al., 2020b), and are characterized by the presence of serine-threonine protein kinase catalytic domain (Wang et al., 2015). As the name indicates, TSSK genes are predominantly expressed in testis and are conserved across diverse species. Although the homologues of TSSK genes have been identified in various organisms including mammals (Kueng et al., 1997), birds (Słowińska et al., 2020), molluscs (Kim et al., 2019) and some insects (Liu et al., 2024; Zhai et al., 2023), their presence and function remains unexplored in mosquitoes.

The present study identified TSSK homologues*, As_tssk3* and *Aea_tssk1* in two mosquito species, *Anopheles stephensi* and *Aedes aegypti*, respectively. Consistent with the findings in other organisms, both *As_tssk3* and *Aea_tssk1* exhibit a male-specific expression, with significantly higher expression of *As_tssk3* in the testes of *An. stephensi* (Fig. 2 and S1). Feeding *An. stephensi* larvae with dsRNA targeting *As_tssk3* led to a significant reduction in its expression in male pupae and adults (Fig. 3 C, D). This once again confirms the effectiveness of droplet feeding method for dsRNA delivery at the larval stage in mosquitoes (Dhandapani et al., 2020). The effect of this knockdown was most notable in the size of the sperm reservoir, resulting in significantly smaller sperm reservoir in knockdown males as compared to that in control males (Fig. 4). Sperm reservoirs are specialised structures in insects that stores matured sperms until it is transferred to female at the time of mating (Huho et al., 2006; Manas et al., 2024). Studies in some insects have correlated the size of sperm reservoir with the number and quality of sperm produced. A smaller sperm reservoir is an indicator of reduced sperm count and an inferior quality of sperm, ultimately affecting male fertility (El-sabrout et al., 2021; Sales et al., 2018). Another significant finding of our study is the reduction in the expression of *protamine* (*As_prot*) and ‘*mosquito testes-specific*’ (*As_mts*) genes in *As_tssk3* knockdown males (Fig. 3 C, 3D). *Protamine* is known to play crucial roles in chromatin dynamics during spermatogenesis and male fertility in insects and other organisms (Balhorn, 2007; Leyden et al., 2024). The histone replacement by protamines in sperm chromatin helps in sperm DNA compaction, an important aspect of spermiogenesis (Zhang et al., 2023). While the *mts* gene is identified as sperm-specific marker that can be used as control marker for sperm numbers also (Krzywinska & Krzywinski, 2009).

Despite a smaller size of sperm reservoir and reduced expression of *As_prot* and *As_mts* genes in *As_tssk3* knockdown males, they were equally fertile as that control males. Females mated with knockdown males laid almost equal number of eggs as those mated with control males (Table S4). At present, the specific reason behind this unexpected observation is unclear, however, one plausible explanation is that mosquitoes possess the remarkable capacity of rapid and continuous sperm production (Damiens et al., 2013; Riemann et al., 1974), which may mitigate the adverse effects of reduced sperm reservoir. Alternatively, the normal fertility levels of *As_tssk3* knockdown males could also be due to alternate adjustment of sperm allocation in these males (Nayyab et al., 2021), as seen in other insects. Furthermore, studies have demonstrated that the fecundity of female mosquitoes doesn’t get compromised if they receive a threshold quantity of sperm from males (De Jesus & Reiskind, 2016; Helinski & Harrington, 2011), suggesting that the reduction in the size of sperm reservoir in knockdown males might not have decreased the sperm production to an extent which would affects the female fecundity.

An alternate hypothesis comes from the compensatory mechanisms in other genes or pathways involved in spermatogenesis which may have maintained the quality and quantity of the sperms in the knockdown males (Cedden & Bucher, 2024; Y. Li et al., 2024; Yuan et al., 2015; Zuo et al., 2022). The above hypothesized mechanisms, individually or in combination, would have maintained the fertility of *As_tssk3* knockdown males and unaffected egg laying pattern in control and target females in our experiment.

Our findings indicate that *As_prot* and *As_mts* genes function downstream to *As_tssk3*, where *As_tssk3* likely regulates their expression via a transcription factor. To explore this potential transcriptional regulator, we analysed the targets of *Drosophila* TSSK (*dTSSK*) related to spermatogenesis process. This analysis indicated ‘Nucleosome Assembly Protein’ (NAP) as the likely candidate (Zhang et al., 2023). We searched for the homologue (As_NAP1) of *Drosophila* NAP in *An. stephensi* and confirmed the presence of multiple phosphorylation sites in the As_NAP1. This indicated As_NAP1 as the likely target of As_TSSK3. This hypothesis was further supported by the strong interaction observed between As_TSSK3 and As_NAP1 in our in-silico interaction studies. In-vitro binding experiments will further establish the link between *As_tssk3*, As_NAP1 and As_Prot.

In summary, this manuscript presents the first identification of TSSK homologue and its potential role in spermatogenesis within mosquito species which further expands our knowledge of TSSKs in non-model organisms. Future studies using genome manipulation to silence *As_tssk3* and its homologues in *Anopheles* and other mosquito species will further shed light on their roles in spermatogenesis, presenting promising applications for mosquito population control. This research could also enhance sterile insect technology, offering a targeted approach for reducing mosquito populations, in future.

## AUTHOR CONTRIBUTIONS

KKM: Writing– original draft; writing– review and editing; conceptualization; investigation; methodology; data curation; validation; formal analysis. CC: Writing– review and editing; methodology. JNS: Supervision; project administration; funding acquisition; writing– review and editing; resources; conceptualization. PR: Mosquito culture maintenance, tissue dissection and real-time PCR. RD: Provided the *Anopheles stephensi* cultures. VS: Provided RNA samples of *Aedes aegypti* developmental stages. All authors contributed to the article and approved the submitted version. KKM is a PhD student at Central University of Rajasthan and this manuscript is a part of his PhD work.

## ACKNOWLEDGEMENTS

JNS acknowledges funding and support from Department of Science and Technology, Ministry of Science and Technology, India, for the EMR and CRG grants (CRG/2023/006907, EMR/2017/001378). We are thankful for the qRT-PCR facility of the DBT Builder project (BT/INF/22/SP44383/2021).

